# Sigma Ligand-mediated Cytosolic Calcium Modulation in BV2 Microglia Cells

**DOI:** 10.1101/2024.11.18.624024

**Authors:** Eric Lodholz, Faria Anjum Simin, Michael Crider, William Neumann, Joseph Schober

## Abstract

While initially protective, prolonged neuroinflammation can lead to a disrupted brain microenvironment and neurodegenerative diseases. Microglia cells are the primary regulators of neuroinflammation, with anti- or pro-inflammatory phenotypes. Just as microglia are crucial in neuroinflammation, calcium is a crucial regulator in the activation of the microglia. Intracellular calcium dynamics can be modulated by small, membrane bound chaperone proteins called sigma receptors. Both sigma receptors 1 and 2, which are involved in a wide variety of cellular functions, have been implicated in neurodegenerative diseases, psychiatry, and cancer. We examined the use of sigma-receptor ligands to modulate cytosolic calcium levels in BV2 cells, an immortalized mouse microglia cell line. Immunofluorescence staining detected both sigma 1 and 2 receptors in perinuclear regions and the cell cytoplasm. Our selection of compounds was a mixture of commercially available sigma receptor ligands and compounds synthesized at Southern Illinois University Edwardsville. We used the Fluo-8-AM calcium probe to measure cytosolic calcium concentrations using flow cytometry after 15-, 25- and 35-minute ligand exposure. With 1 µM ligand concentration, we found significantly increased cytosolic calcium levels in BV2 cells after 15-minute exposure. A moderate correlation was determined between sigma receptor 2 selectivity and calcium activity, suggesting a sigma-mediated calcium effect. In addition, we aimed to determine the location of calcium flux by pretreating cells with thapsigargin and EGTA to inhibit both the endoplasmic reticulum and extracellular space of suspected calcium entry. We found all four compounds tested, siramesine, PB-28, cis-8-OMe BBZI, and BN-IX-111-F1 promote calcium entry from the extracellular space, but only siramesine promotes calcium entry from both locations. We tested for a common drug-induced effect called phospholipidosis using HCS LipidTOX phospholipid detection reagent to determine a correlation between calcium activity and a membrane-mediated effect. Only two out of the five compounds tested, PB-28 and amiodarone, resulted in significant phospholipid accumulation at 10 µM treatment, suggesting little to no correlation between calcium levels and phospholipidosis. Further investigations are required to target the exact mechanism of calcium flux and characterize the connection between phospholipidosis and sigma receptors after sigma ligand exposure. Nevertheless, our work emphasizes the importance of sigma ligand-modulated intracellular calcium dynamics as a potential route of therapy, specifically for modulation of neuroinflammation.

## Introduction

Calcium is one of the most important second messengers and is involved in many, if not all, aspects of cellular physiology in some way. Calcium signaling facilitates processes like cell proliferation, metabolism, motility, protein phosphorylation, gene transcription, and many others (Bootman and Bultynck, 2020; Missiroli et al., 2020). Intracellular calcium signaling is regulated through ion channels, active pumps, voltage-gated channels, ion exchangers, and store-operated calcium entry. Calcium flux is also active in and out of intracellular stores like the ER, mitochondria, and lysosomes (Patergnani et al., 2020). Because of its involvement in many cellular functions, alterations in calcium regulation can lead to a wide array of physiological dysfunctions and pathologies like neurodegenerative diseases, cancer, diabetes, and more (Pontisso and Combettes, 2021). Microglia cells are electrically non-excitable, so the primary route for calcium signaling is through intracellular stores, ligand-gated ion channels on the plasma membrane, and store-operated calcium entry (Kettenmann et al., 2011).

Sigma receptors (SR) are multi-functioning proteins enriched in the CNS and expressed by microglia (Jia et al., 2018). They are classified into two subtypes: SR1 and SR2. SR1 has no genetic relationship to any other human gene and is genetically unrelated to SR2 (Alon et al., 2021; Schmidt and Kruse, 2019). Most ligand binding occurs on the hydrophobic C-terminus domain on the cytosolic side, mainly due to an electrostatic interaction between Glu172 and a basic nitrogen (Schmidt and Kruse, 2019; Schmidt et al., 2016). SR1 can bind a large variety of structurally diverse ligands (Walker et al., 1990). The current pharmacophore model of SR1 is a central basic nitrogen flanked by two hydrophobic/aromatic moieties that are not equidistant from the center (Fallica et al., 2021; Szczepańska et al., 2022). This pharmacophore is very similar to the SR2 pharmacophore, indicating an overlap in structural preference for both receptors (Abate et al., 2009; Szczepańska et al., 2022).

SR1 has been characterized as a molecular chaperone protein that modulates many cellular signaling pathways including GPCRs, ion channels, and IP3R (Schmidt and Kruse, 2019; Hayashi and Su, 2007). Therapeutic potential of SR1 modulation includes cellular functions involved in cancer growth, psychiatric disorders, pain management, stroke, ALS, Parkinson’s, and Alzheimer’s disease (Romero et al., 2016; Nguyen et al., 2015; Hayashi, 2019). SR1 is localized in the ER membrane and to lesser extent in the plasma membrane where it can modulate different ion channels and receptors (Hayashi and Su, 2007; Nguyen et al., 2017). SR1 modulates calcium flux between the ER and mitochondria at mitochondria-associated membranes (MAM) by stabilizing IP3R channel forming complexes (Hayashi and Su, 2007; Ooi et al., 2021). IP3R can be quickly degraded via ubiquitination after IP3 binds in the absence of SR1 chaperone activity (Alzayady and Wojcikiewicz, 2005). Studies have also shown the addition of SR1 ligands activate SR1 and stabilize the IP3R-GRP75-VDAC complex, subsequently rescuing calcium signaling at the MAM and even promoting a neuroprotective microglial activation route (Ooi et al., 2021; Nguyen et al., 2017). With this mechanism, SR1 may support a neuroprotective feature by facilitating calcium flow to mitochondria across a MAM. SR1 may induce cytosolic calcium increase from the ER at the IP3R. In response to ER-calcium depletion, STIM can bind to SR1 at the PM-ER membrane junction, facilitating calcium release via the SOCE mechanism (Herrando-Grabulosa et al., 2021). SR1 has also been shown to regulate calcium influx after translocation to the PM, interacting with ion channels and GPCRs to potentially increase cytosolic calcium levels (Schmidt and Kruse, 2019). In contrast, SR1 activation can also inhibit calcium influx through voltage-gated calcium channels in retinal ganglion cells (RGCs) (Mueller et al., 2013; Tchedre et al., 2008).

SR2 was first identified as the progesterone membrane binding component-1 (PGRMC1) based on radioligand binding experiments of SR2 ligands and knockdown and overexpression experiments (Xu et al., 2011). This identification was debated, as later studies revealed SR2 and PGRMC1 were two different proteins (Pati et al., 2017). It is now widely accepted that the SR2 identity is an ER-resident transmembrane protein called TMEM97 involved in cholesterol homeostasis and Niemann-Pick type C disease (Alon et al., 2017). However, little is known about the SR2 cellular and molecular functions. Another study proposed its involvement with low-density lipoprotein (LDL), leading to increased LDL internalization (Riad et al., 2018). SR2 is also highly expressed in the CNS and peripheral tissues like the liver and kidney (Alon et al., 2017; Alon et al., 2021). Two SR2 ligands are currently in clinical development for their therapeutic potential in Alzheimer’s and schizophrenia (Grundman et al., 2019; Harvey et al., 2020). Additionally, SR2 is highly expressed in cancer cells, prompting its use as a cancer biomarker and in studies for its therapeutic potential (van Waarde et al., 2015; Zeng et al., 2020). In terms of calcium regulation, a few studies propose SR2’s ability to modulate cytosolic calcium levels upon ligand stimulation, inducing both transient and non-transient increases (Vilner and Bowen, 2000; Cassano et al., 2009). With SR2’s localization at the ER membrane, its calcium regulation abilities are reasonable.

Given the evidence of the SR effect on calcium dynamics, we proposed the use of SR ligands on BV2 cells to modulate cytosolic calcium levels, thereby influencing microglial activation. We treated samples of BV2 cells with 18 different ligands and measured their resulting cytosolic calcium levels using flow cytometry. We determined the location of the calcium through pretreatments with thapsigargin and EGTA to inhibit ER and extracellular sources of calcium, respectively. We tested select compounds from the calcium assay for their induction of phospholipidosis, a common drug-induced side effect involving phospholipid accumulation and lysosomal dysfunction (Breiden and Sandhoff, 2020). In short, we investigated a potential alternative mechanism for the calcium effect, focusing on a membrane-mediated route leading to phospholipidosis, rather than a SR-mediated route causing direct calcium level changes. We hypothesized there would be a correlation between the calcium effect and the propensity for phospholipidosis induction of those same compounds.

## Materials and Methods

### Cell Line, Chemicals, and Reagents

Dulbecco’s Modified Eagle’s Medium (DMEM), phosphate buffered saline (PBS), and fetal bovine serum (FBS) were purchased from Corning Life Sciences (Manassas, VA). Bovine plasma derived fibronectin was obtained from R & D Systems (Minneapolis, MN). Bovine serum albumin (BSA) was purchased from GE Healthcare Life Sciences (Chicago, IL). Paraformaldehyde was purchased from Electron Microscopy Sciences (Hatfield, PA). Trypsin-EDTA (0.25%) phenol red was obtained from Thermo Fisher Scientific (Waltham, MA). Aqua-Poly/Mount was purchased from Polysciences (Warrington, PA). Sigma 1 Receptor mouse monoclonal antibody was obtained from Santa Cruz Biotechnology (Dallas, TX). Goat anti-mouse Alexa Fluor™ Plus 488, TMEM97 rabbit polyclonal antibody, and goat anti-rabbit Alexa Fluor™ Plus 647, Hoechst 33258 DNA stain, and Alexa Fluor™ Texas Red phalloidin were purchased from Thermo Fisher Scientific Inc (Waltham, MA). The Fluo-8-AM calcium probe was obtained from AAT Bioquest (Sunnyvale, CA). Triton X-100, EGTA, and Dimethylsulfoxide (DMSO) were obtained from Sigma Aldrich (St. Louis, MO). Thapsigargin was obtained from Tocris (Bristol, UK). The HCS LipidTOX Green Phospholipid Detection Reagent was obtained from Fisher Scientific (Waltham, MA). The Ki determinations for the SIUE compounds (synthesized by Dr. Michael Crider and Dr. Bill Neumann) were provided by the National Institute of Mental Health’s Psychoactive Drug Screening Program, Contract no. HHSN-271-2913-00017-C (NIMH PDSP). The NIMH PDSP is directed by Bryan L. Roth MD, PhD at the University of North Carolina at Chapel Hill and Project Officer Jamie Driscoll at NIMH, Bethesda, MD, USA.

**Table 1:** Compounds used in this study.

| Name | Source | SR1 Ki | SR2 Ki | Ki Source |
| --- | --- | --- | --- | --- |
| AMC-296-39-2 | SIUE | 2.8 | 23 | NIMH PDSP |
| AMC-610 | SIUE | NA | NA | NA |
| AMC-S2-77-2 | SIUE | NA | NA | NA |
| Amiodarone | Thermo Fisher Scientific | 791.7 | >10000 | NIMH PDSP |
| BD-1047 | Tocris Bioscience | 0.93 | 47 | (Matsumoto et al. 1995) |
| BN-IX-111-F1 | SIUE | 463 | 6 | NIMH PDSP |
| Cis-8-OH BBZI | SIUE | 3.6 | 2.2 | NIMH PDSP |
| Cis-8-OH MPZI | SIUE | NA | NA | NA |
| Cis-8-OH PBZI | SIUE | 11 | 6.9 | NIMH PDSP |
| Cis-8-OMe BBZI | SIUE | 0.8 | 2.2 | NIMH PDSP |
| Desipramine | Tocris Bioscience | 1987 | 34630 | (Narita et al. 1996) |
| MD-19A-13-1 | SIUE | 7 | 462 | NIMH PDSP |
| PB-28 | Tocris Bioscience | 0.38 | 0.68 | (Abate et al. 2020, 28) |
| PRE-084 | Sigma-Aldrich | 2.2 | 13091 | (Maurice et al. 1999) |
| Siramesine | Sigma-Aldrich | 17 | 0.12 | (Perregaard et al. 1995) |
| SK&F 10047 | Tocris Bioscience | 1800-4657 | 4500-13694 | (Chou et al. 1999) |
| SM-21 | Tocris Bioscience | >1000 | 67.5 | (Ghelardini, Galeotti, and Bartolini 2000) |
| Trans-8-OH ABZI | SIUE | 15 | 11 | NIMH PDSP |
| SIUE- Compounds were synthesized in the Department of Pharmaceutical Sciences at Southern Illinois University Edwardsville<br>Ki- Units are given in nanomolar<br>NA- Structures are based on other compounds with known Ki values.<br>NIMH PDSP- Ki values were obtained from the NIHMH Psychoactive Drug Screening Program from the Roth Lab at University of North Carolina at Chapel Hill. |  |  |  |  |

### BV2 Microglia Cell Culture

BV2 microglia cells were generated from the retrovirus immortalization of mouse microglia cells because of their preservation of primary mouse-microglia phenotypes (Blasi et al. 1990). Cells were maintained in 25 cm^2^ flasks with 5 mL growth media (DMEM, 10% FBS and penicillin-streptomycin-amphotericin B) in a 37°C and 5% CO_2_ and passaged after they reached 80-90% confluency. Cells were trypsinized and suspended in growth medium. Cell suspension density was adjusted to 200 cells/μL for epifluorescence microscopy and 100 cells/μL for calcium assay.

### Epifluorescence Microscopy

Glass coverslips were coated with 60 μg/mL fibronectin in PBS for an hour at 37°C to ensure adequate cell adhesion. The fibronectin-coated coverslips were placed in 12-well plates and 1 mL of cell suspension was added and incubated for 3 hours at 37°C, 5% CO_2_. The cells were then fixed overnight in 4% paraformaldehyde and 0.1% Triton X-100 in PBS and after fixation, they were washed in deionized water. The cells were blocked with 1% BSA in PBS for 20 min at 37° C and were incubated with Sigma 1 Receptor mouse monoclonal antibody and TMEM97 (sigma 2 receptor) rabbit polyclonal antibody for 20 min at 37° C. The coverslips were washed in deionized water and placed 0.1% BSA for 10 mins. The cells were incubated with Hoechst 33258 (DNA/nuclear stain), Alexa Fluor Texas red phalloidin (actin stain), goat anti-mouse Alexa Fluor 488, and goat anti-rabbit Alexa Fluor 647 secondary antibodies for 20 min at 37° C. The coverslips were washed in deionized water, placed in PBS for 10 min, and washed in deionized water again. Lastly, the coverslips were mounted on glass slide using Aqua-Poly/Mount. The images were acquired at 63X oil immersion objective using a Leica DMi8 inverted microscope (Leica, Wetzlar, Germany) fitted with 12-bit charge-coupled device monochrome camera.

### Image Processing and Analysis

The acquired images were processed and analyzed using MetaMorph Microscopy Automation and Image Analysis Software. The images were adjusted for uneven intensity using a size-based background estimation process and de-noised with a low pass filter followed by subtraction of the appropriate lower threshold values. Then the inclusive threshold was applied to select objects of interest for the analysis. 30 cells were chosen randomly and subjected to quantitative analysis for sigma 1 and sigma 2 receptors. Actin staining was used to obtain the pixel area of individual cells. The size and number of distinct objects in pixels was recorded with Integrated Morphometry Analysis tool and the average was calculated for each cell for sigma 1 and sigma 2 receptor. The pixel values were converted to micron squared by calibration using a stage calibrator slide (1 pixel = 0.0196 µ² in 63X objective). Then colocalization of sigma 1 receptor and sigma 2 receptor was recorded by means of percent overlap using following formula,

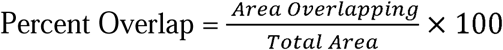

### Fluo-8-AM Probe Cytosolic Calcium Assay of Sigma Ligands

The Triton X-100 concentration that resulted in the maximum Fluo-8-AM calcium probe fluorescence was determined and the max fluorescence was used to calculate the real concentration of calcium in the cell cytosol. Three concentrations of each ligand were tested (1µM, 100 nM, and 10 nM). Each experiment also included one sample with 0.01% Triton X-100, one 0.03% DMSO control, and one growth medium only control. 2000 nM of Fluo-8-AM calcium probe was added to cell suspensions and incubated for 30 minutes at 37°C. Fluorescence was measured at 520 nm wavelength using the Accuri C6 flow cytometer (BD Biosciences, San Jose, CA, USA) at three timepoints: 15 minutes, 25 minutes, and 35 minutes post-addition of ligand.

### Thapisgargin/EGTA Assay

Four sigma ligands with most calcium activity (15 min, 1µM) from the cytosolic assay were tested for origin of calcium ion flux post exposure. Thapsigargin pretreatment was used to deplete endoplasmic reticulum calcium and EGTA pretreatment was used to chelate extracellular calcium. Cell suspensions were first incubated with Fluo-8-AM probe for 30 min at 37°C and then pretreated with either 1 µM Thapsigargin or, 1 mM EGTA for 10 min. After pretreatment, 1 µM sigma receptor ligand was added and incubated for an additional 15 min at 37°C. This assay also included ligand treatment with no Thapsigargin and EGTA pretreatment, 0.01% Triton X-100, and growth medium only control. After 15 min fluorescence was measured using the flow cytometer.

### Phospholipidosis Assay

The HCS LipidTOX Green Phospholipidosis Detection Reagent probe was used to determine phospholipid accumulation in BV2 cells after drug exposure. Five sigma ligands were tested in this assay: Amiodarone, SK&F 10047, PRE-084, Cis-8-OMe BBZI, and PB-28. BV2 cells were treated with 0.1, 1.0, and 10 µM ligands and DMSO control for 24 hours. After the 24-hour treatment, LipidTOX probe was added, and the cells were then incubated at 37°C for 24 hours. The cells were trypsinized and then transferred to centrifuge tubes with 0.5 mL growth media to allow measurement on the flow cytometer.

### Data Analysis

The raw fluorescence data for each sample per ligand was pooled together using GraphPad Prism to provide an average fluorescence. The average fluorescence data was then converted into nanomolar calcium concentration using the Kd value of the probe provided by th manufacturer according to the following relationship,

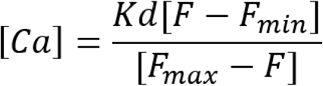

where the Kd reported by the manufacturer is 389 nM, *F* is the experimental fluorescence, is the auto-fluorescence of a BV2 cells, is the maximum fluorescence after maximum cytosolic calcium entry from outside the cell after 0.01% Tx100 treatment. The resulting nanomolar calcium values in each time group (15 min, 25 min, 35 min) were normalized to th 15-minute control mean of each experiment.

### Statistical Analysis

Using GraphPad Prism, a one-way analysis of variance (ANOVA) test and Dunnett’s Multiple Comparison’s test were performed on all graphs. The specific comparisons within samples of each assay varied and are denoted in their respective section. The notation of significance is as follows: *** = P value < 0.001, ** = P value 0.001 to 0.01, * = P value 0.01 to 0.05, and ns = not significant (P value > 0.05).

### Cytosolic [Ca^2+^] and Sigma Receptor Selectivity

Correlations were examined by performing linear regression analysis on GraphPad Prism to determine the coefficient of determination ( ). Sigma 2 receptor selectivity was determined by dividing sigma 1 receptor Ki by the sigma 2 receptor Ki. The Ki data for SIUE compounds were provided by the National Institute of Mental Health’s Psychoactive Drug Screening Program.

## Results

### Sigma receptor intracellular localization and colocalization

Sigma receptors are primarily localized in cytoplasm of BV2 cells (**Figure 1**). On average, sigma 1 receptors account for 10.1 % and sigma 2 receptors account for 7.4% of the total cell area. 16.7% of total area of sigma 1 receptors colocalize with sigma 2 receptors and 22.7 % of total area of sigma 2 receptors colocalize with sigma 1 receptors. There is no monotonic relationship between relative intensities where sigma receptors colocalize.

**Figure 1.**
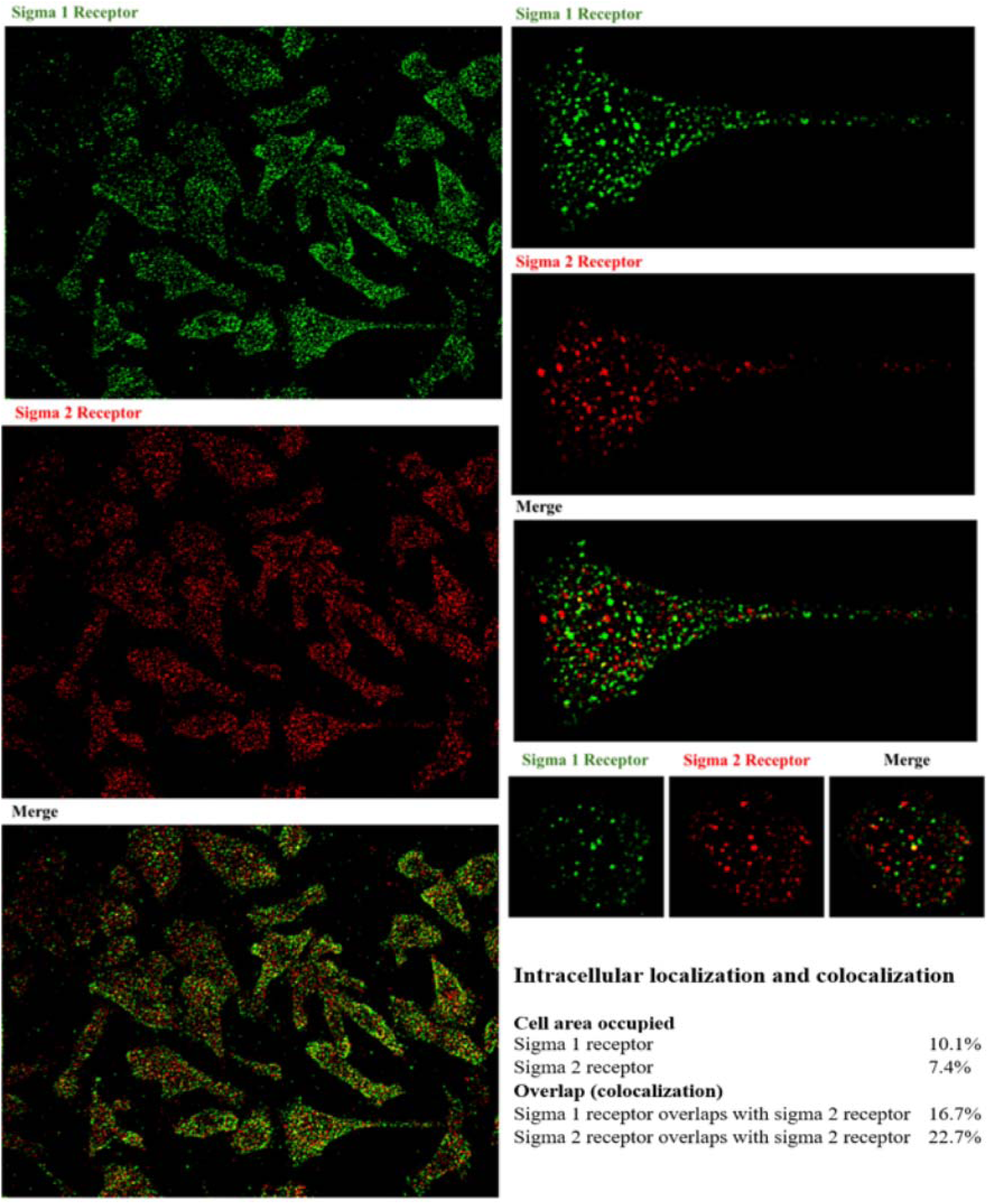
Expression of sigma receptors in BV2 cells. BV2 cells were fixed overnight and then stained for sigma receptors, nucleus, and actin. Images were processed to quantify sigma receptor colocalization.

### Cytosolic Calcium Modulation After Ligand Exposure to BV2 Cells

At 1 µM concentration (**Figure 2**), AMC-296-39-2, AMC-610, AMC-S2-77-2, Amiodarone, BD-1047, BN-IX-111, Cis-8-OH MPZI, Cis-8-OMe BBZI, MD-19A-13-1, PB-28, PRE-084, Siramesine, SK&F 10047, and SM-21 tested with significantly increased cytosolic calcium levels in BV2 microglia cells after 15-minute exposure. Siramesine showed increased [Ca^2+^] at 100 nM concentration after 15-, 25-, and 35-minute exposure. No significant increase in cytosolic [Ca^2+^] was observed with all ligands at 10 nM concentration at all timepoint. Cis-8-OMe-BBZI and PB-28 increased cytosolic calcium levels at 100 nM concentration after 25 minutes.

**Figure 2.**
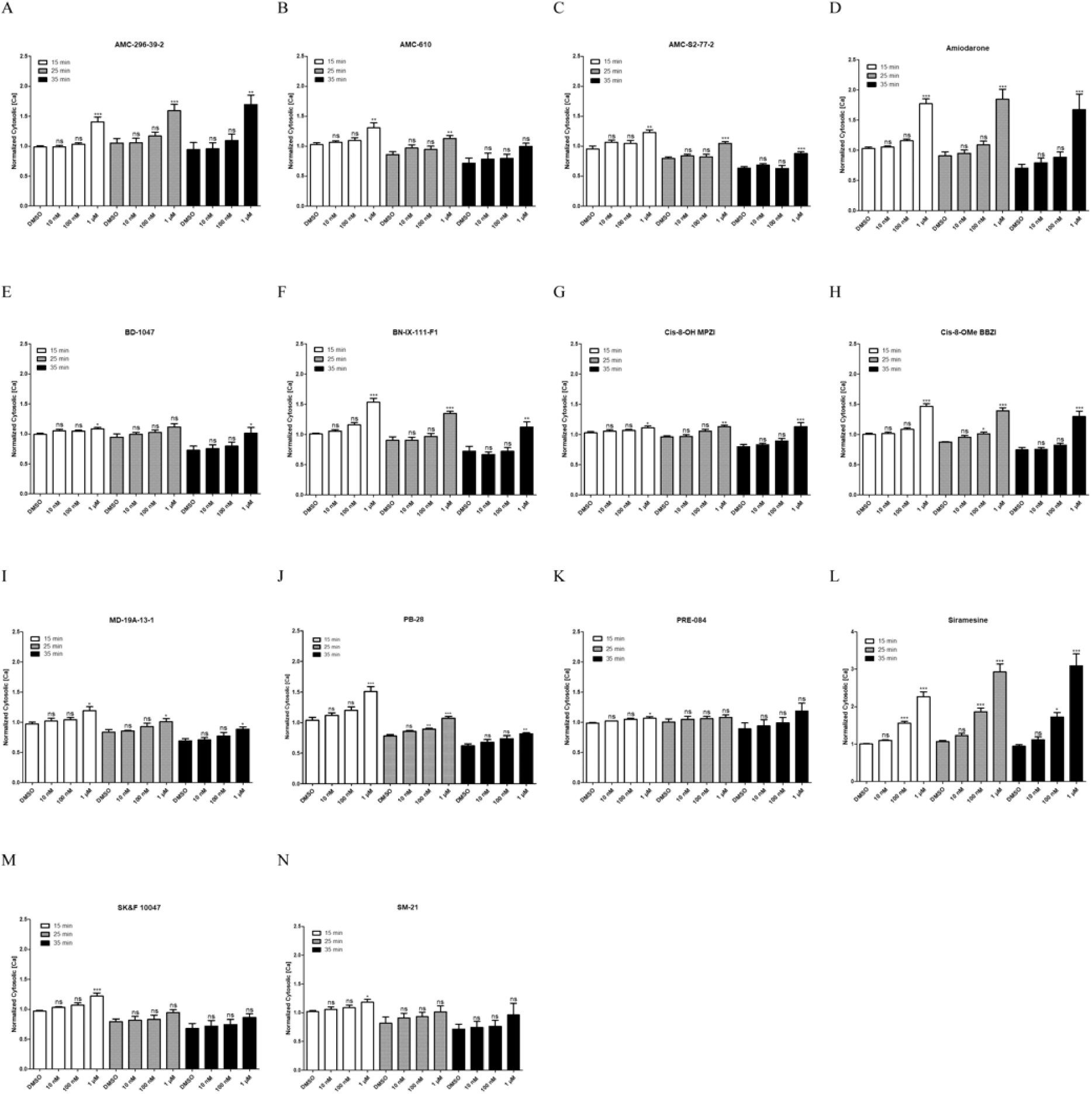
Sigma ligands effects on cytosolic [Ca^2+^] at 15, 25 and 35 minutes. At 1 µM concentration, (A) AMC-296-39-2, (B) AMC-610, (C) AMC-S2-77-2, (D) Amiodarone, (E) BD-1047, (F) BN-IX-111, (G) Cis-8-OH MPZI, (H) Cis-8-OMe BBZI, (I) MD-19A-13-1, (J) PB-28, (K) PRE-084, (L) Siramesine, (M) SK&F 10047, and (N) SM-21 caused significant increase in cytosolic [Ca^2+^] after 15 minutes. All data were normalized to the 15-minute control and a one-way analysis of variance (ANOVA) test and Dunnett’s Multiple Comparison’s test were performed.

### Ranking of all Ligands by Cytosolic [Ca^2+^] Levels over 35 Minutes at 1 µM and 100 nM Concentration

Siramesine had the highest potency at all three time points. The remaining order of compounds varied for each time point, indicating a wide range of declining rates over time (**Figure 3**). The least active compound at 1 µM for 15, 25, and 35 minutes was PRE-084, SK&F 10047, and PB-28, respectively. For 100 nM, the least active compound at 15 minutes was Cis-8-OH PBZI and at 25 and 35 minutes it was AMC-S2-77-2. Compared to the 1 µM rankings, the later time points for 100 nM treatments began to result in more [Ca^2+^] values well below the 15-minute control mean, illustrating the significance drug concentration has on resulting cytosolic [Ca^2+^] activity. All graphs were normalized to the 15-minute control which is indicated by 1 on the x-axis.

**Figure 3.**
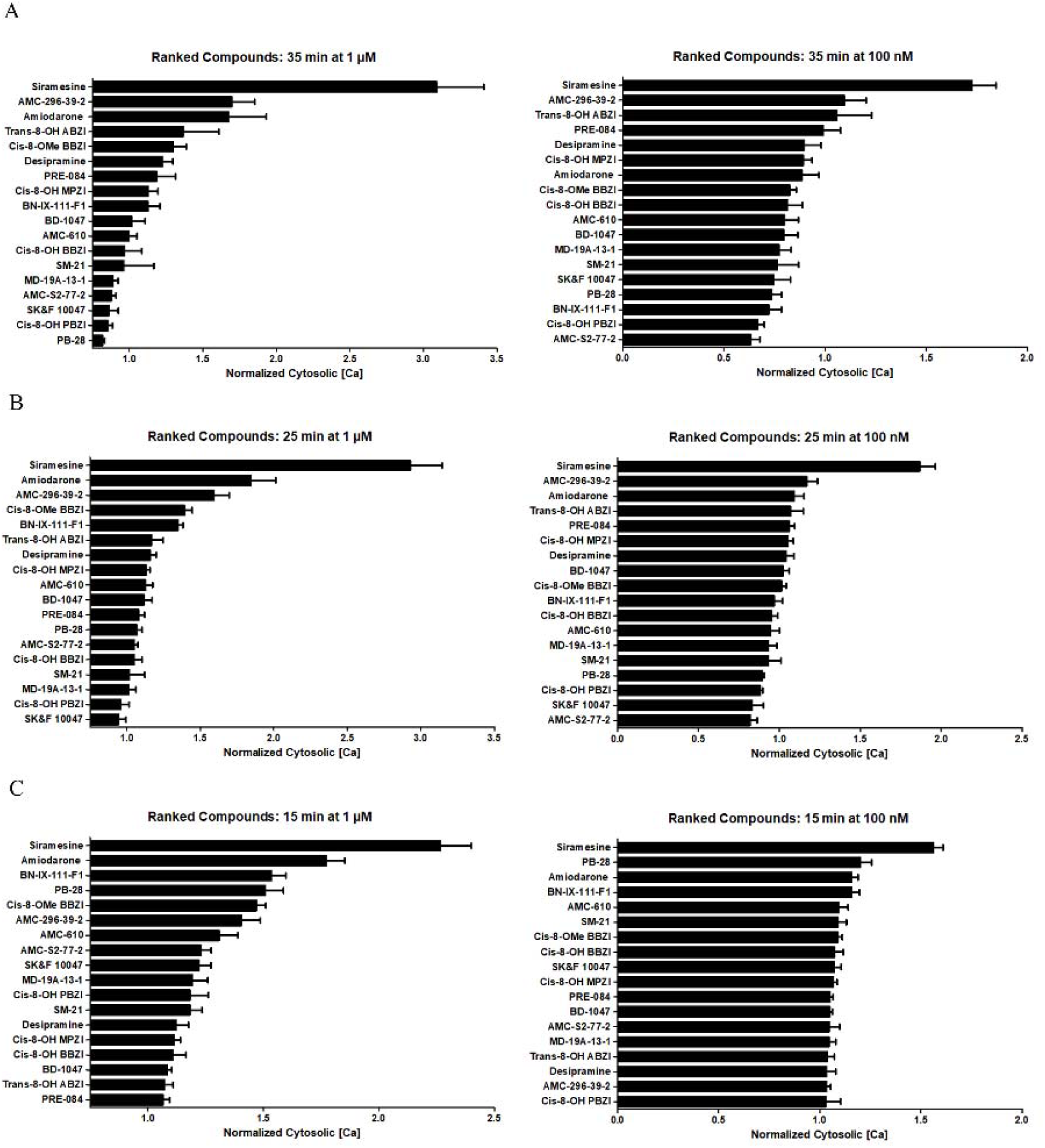
The ranking of all compounds cytosolic [Ca^2+^] levels at 1 µM and 100 nm concentration after (A) 35 minutes, (B) 25 minutes, and (C) 15 minutes. All data were normalized to the 15-minute control which is indicated by 1 on the x-axis.

### Correlation Between Cytosolic [Ca^2+^] and Sigma Receptor Ki

The normalized [Ca^2+^] levels after 15 minutes of 1 µM ligand exposure were graphed against their respective sigma 1 and 2 receptor Ki values. There was no correlation when [Ca^2+^] levels were paired against each ligand’s sigma 1 or 2 receptor Ki after obtaining the coefficient of determination (r^2^) through a linear regression analysis (**Figure 4**). However, a moderate correlation (r^2^) was noted when [Ca^2+^] levels were paired against sigma 2 receptor selectivity of each ligand (**Figure 5**). Siramesine was the most sigma 2 receptor selective compound and provided the largest [Ca^2+^] increase. AMC-S2-77-2, AMC-610, and Cis-8-OH MPZI were not included as their Ki data for sigma receptors had not yet been determined, or in the case of Cis-8-OH MPZI, had no sigma receptor affinity.

**Figure 4.**
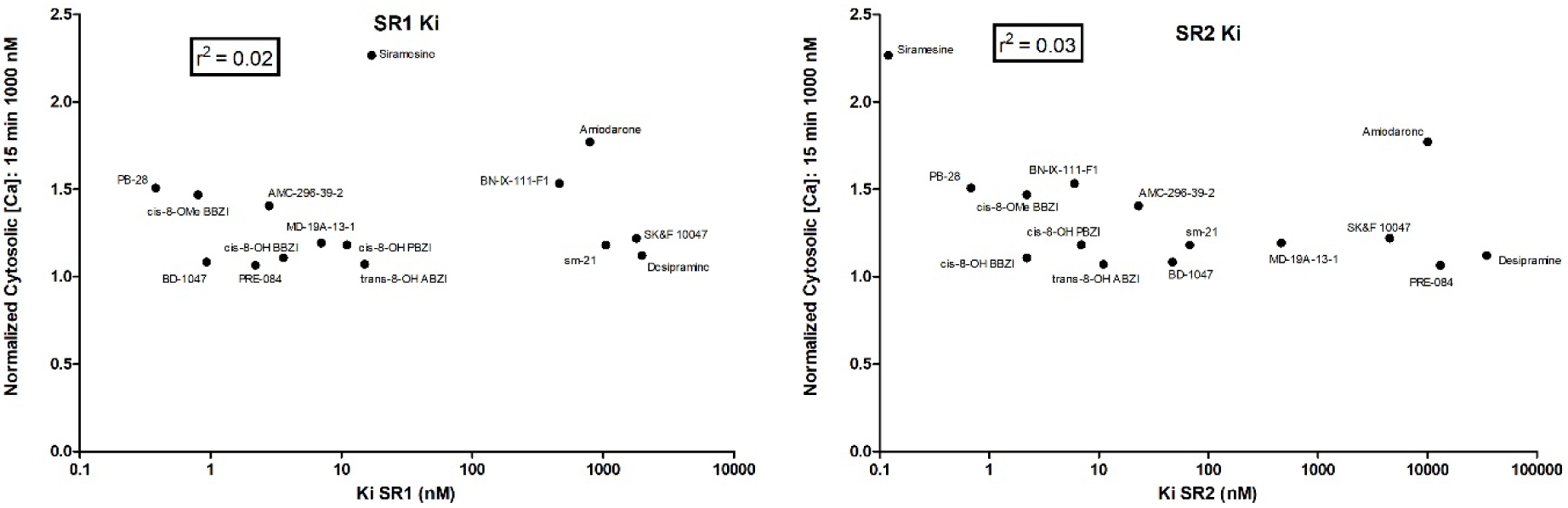
Cytosolic [Ca^2+^] levels after 15 minutes of 1 µM drug exposure paired with sigma 1 receptor Ki and sigma 2 receptor Ki of each drug. There was no correlation when [Ca^2+^] levels were paired against each ligand’s sigma 1 and sigma 2 receptor Ki after obtaining the coefficient of determination (r^2^) through a linear regression analysis.

**Figure 5.**
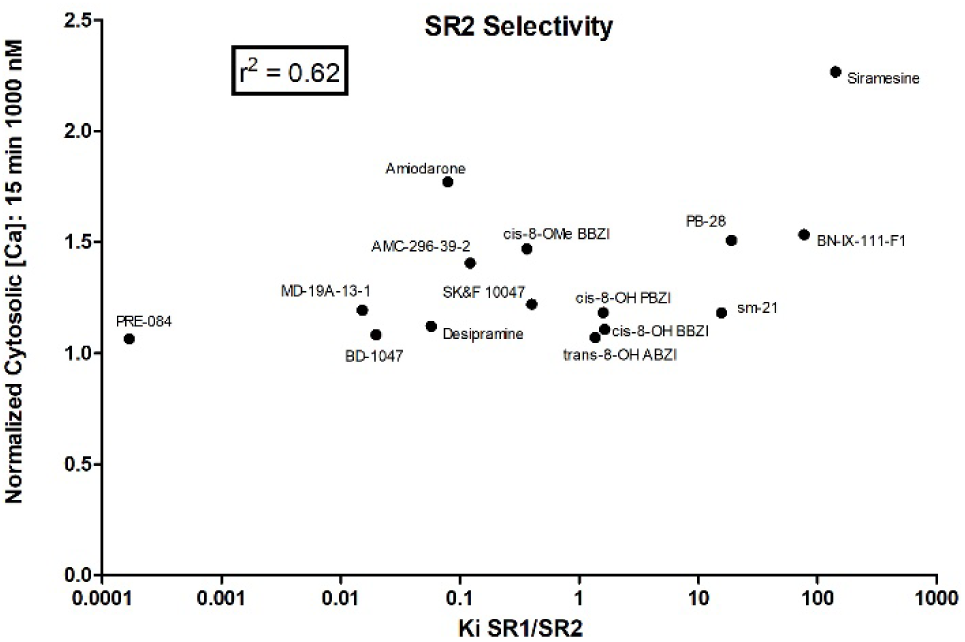
Correlation between Cytosolic [Ca^2+^] after 15 minutes at 1 µM concentration and sigma 2 receptor selectivity. A moderate correlation (r^2^) was noted when [Ca^2+^] levels were paired against sigma receptor selectivity of each ligand.

### Cytosolic Calcium Modulation of Ligands after EGTA and thapsigargin (TG) Pretreatment

In these experiments, we tested the effects of TG and EGTA pretreatment of the sigma-ligand induced increases in cytosolic calcium concentration. Following TG or EGTA, cells were stimulated with 1 µM sigma compound for 15 minutes. For siramesine, both EGTA and TG pretreatment significantly inhibited cytosolic calcium activity (**Figure 6)**. In contrast, for PB-28, Cis-8-OMe BBZI and BN-IX-111-F1, EGTA significantly inhibited cytosolic calcium levels, but not TG.

**Figure 6.**
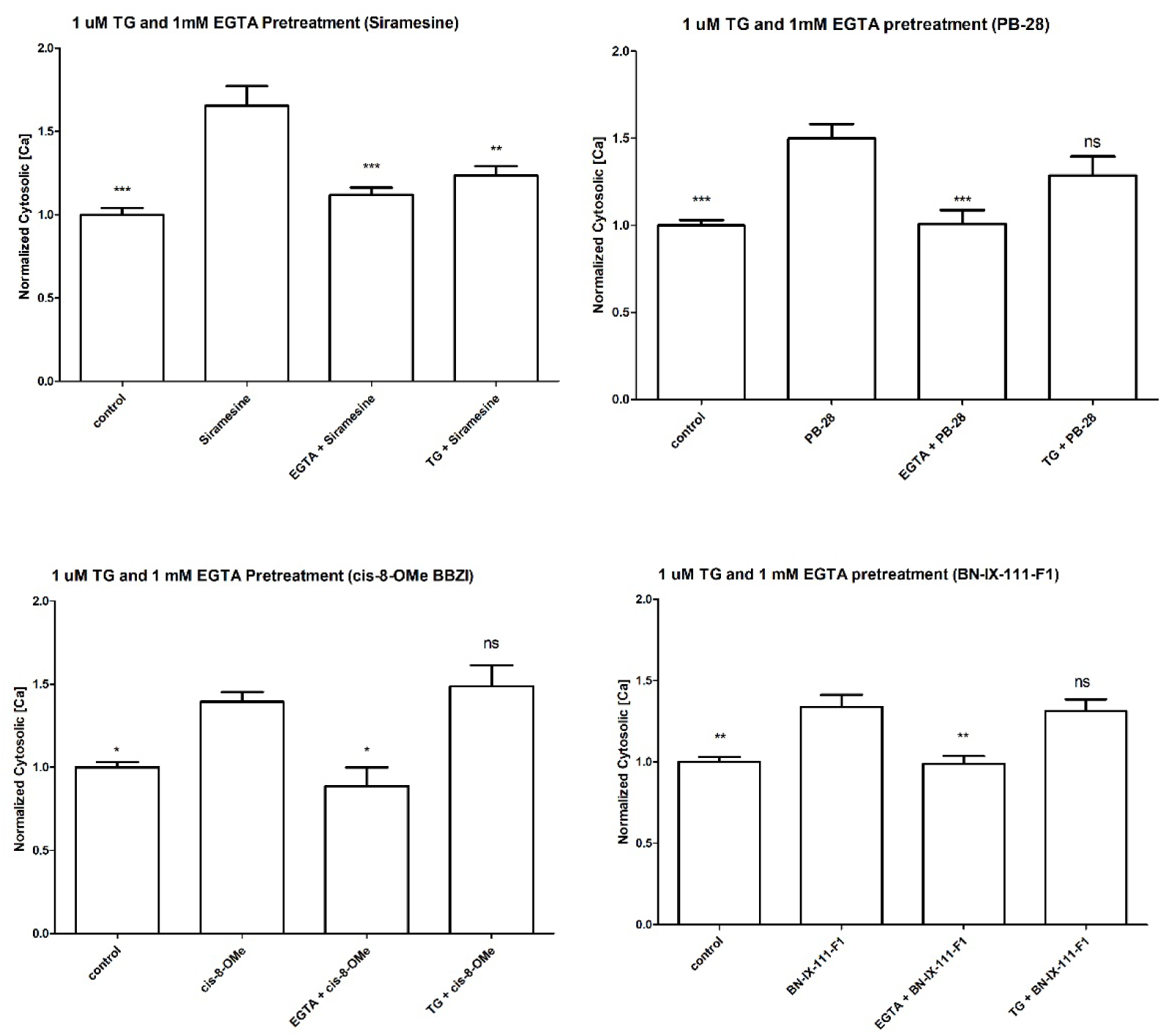
Cytosolic Calcium Modulation of Siramesine, PB-28, Cis-8-OMe BBZI, and BN-IX-111-F1 after EGTA and thapsigargin (TG) pretreatment. The cells were pretreated with 1 mM EGTA or 1 µM TG before addition of 1 µM compound. Cytosolic calcium was measured after 15-minute compound exposure. Statistical comparisons were made against 1 µM compound alone.

### Effect of Sigma Receptor Ligands on Phospholipid Accumulation

In the initial studies, none of the 5 selected compounds (tested at 10 µM) increased phospholipidosis at 15, 25, or 35 minutes (data not shown). Two of the compounds, amiodarone and PB-28, resulted in significant phospholipid accumulation at 10 μM treatment after 24 hours (**Figure 7**). No significant phospholipidosis was observed with SK&F 10047, PRE-084, and Cis-8-OMe BBZI at any of the tested concentrations at 24 hours. However, SK&F decreased phospholipidosis at all three concentrations.

**Figure 7.**
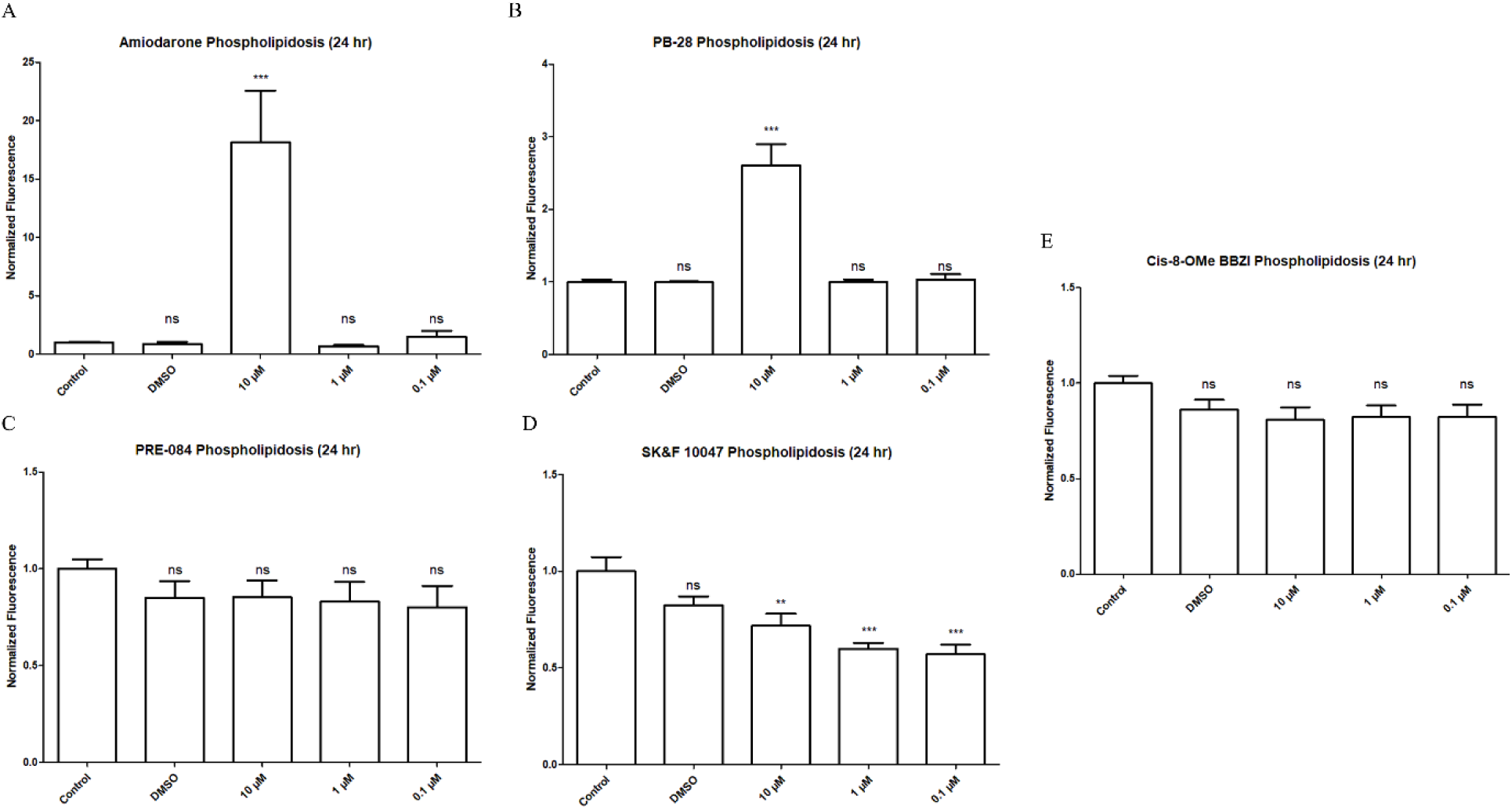
Sigma receptor ligands (A) Amiodarone, (B) PB-28 mediated, (C) SK&F 10047, (D) PRE-084, and (E) Cis-8-OMe BBZI phospholipid accumulation after 24 hours.

## Discussion

This research provided insight into the quantitative subcellular localization and colocalization of sigma receptors and cytosolic calcium mechanics of BV2 microglia cells after stimulation with sigma receptor ligands. The first assay involved testing sigma ligands for their effect on calcium modulation. In general, the 1 µM ligand treatment induced the most significant change from control after 15 minutes. The next assay was necessary to determine if sigma receptor selectivity influenced the potency of calcium activity. The third assay helped indicate the primary location of calcium flux after ligand treatment. Finally, we performed a phospholipidosis assay on five ligands to determine which, if any, promoted phospholipid accumulation.

The exact mechanism of phospholipidosis, a common drug induced side-effect, is not clear, but it is known that substances of cationic-amphiphilic nature can induce it, most notably cationic amphiphilic drugs (CAD) (Breiden and Sandhoff 2020). Interestingly, this general CAD structure is moderately consistent with the sigma receptor pharmacophore: a basic nitrogen flanked by two hydrophobic moieties. This supports the idea of a non-saturable, membrane-mediated mechanism of potential cytosolic calcium activity by the sigma receptor ligands, in which they are acting as CADs in membranes and less so occupying the sigma receptor. When we evaluated the correlation of selected sigma receptor ligands that induced phospholipid accumulation to their respective cytosolic calcium levels, amiodarone and PB-28 induced significant phospholipid accumulation. These two drugs are known phospholipidosis inducers (Tummino et al. 2021). For both drugs, phospholipidosis was observed with 10 µM concentration whereas the significant calcium effect was observed with 1 µM concentration.

Thus, these two drugs show a potential connection between high calcium levels and high phospholipid accumulation. Interestingly, Cis-8-OMe BBZI did not induce any significant phospholipid accumulation in the plasma membrane despite inducing high calcium levels. PRE-084 and SK&F 10047 resulted in a low calcium activity and a mild calcium activity, respectively. The low calcium and low phospholipidosis of PRE-084 is reasonable considering the corresponding low activity of each assay. However, SK&F 10047 was more surprising given its moderate calcium activity with phospholipid levels decreasing lower than control. In terms of amiodarone and PB-28, the high calcium levels and high phospholipid levels indicate a possible connection between the two phenomena. Due to the timing differences of both assays, the observed calcium increase could be upstream of phospholipidosis. Again, this connection is not well understood, but we can postulate ideas based on our findings and current knowledge. It is known that sigma 2 receptors are present in lysosomal membranes and that lysosomes are a major intracellular store for calcium (Schmidt and Kruse 2019). Therefore, PB-28 and amiodarone may be activating sigma 2 receptors on the lysosomal membrane, disrupting lysosomal calcium dynamics and inhibiting normal lipid processing necessary to prevent phospholipid accumulation in the lysosome. Calcium dynamics are crucial for a functioning organelle, so it would not be surprising that a disruption could result in impaired degradation mechanisms used by lysosomes. The same could be applied to PRE-084 due to the little calcium activity. Potentially, not enough calcium activity was induced to promote a strong phospholipidosis observed 24 hours later. Possible explanations for the surprising results of Cis-8-OMe BBZI and SK&F 10047 are more difficult. It may be the targeted sigma receptors for these drugs are localized more on the plasma membrane and/or endoplasmic reticulum, resulting in less calcium release from the lysosomes. It is also possible these ligands represent less CAD character than PB-28 and amiodarone. Amiodarone may be more lipophilic due to more aliphatic hydrocarbons than BBZI and SK&F, despite having the same number of ring structures. Further studies are necessary to compare the lipophilic character of these drugs to amiodarone and PB-28.

We wanted to evaluate a possible correlation between sigma 1 and 2 receptor Ki of each drug and its calcium activity. Based on a low r^2^, it was evident that there was no correlation between both sigma 1 and 2 receptor Ki and calcium activity. We would expect the drugs with higher affinity for sigma receptors might occupy more sigma receptors and subsequently induce higher cytosolic calcium levels. This indicates a possible non-sigma receptor mediated route for the calcium activity. Although individual sigma 1 and 2 receptor Ki did not correlate with calcium levels, there was moderate correlation between sigma 2 receptor selectivity and respective calcium activity. This correlation indicates more of a sigma 2 receptor mediated calcium effect. It is also possible that drugs like amiodarone and PB-28 are inducing calcium release through both phospholipid accumulation and a receptor mediated effect.

Using thapsigargin and EGTA, we wanted to determine the source of calcium entry into the cytosol with the top four active compounds in the calcium assay. There was a significant decrease in calcium levels after pretreatment with EGTA after 15 minutes which indicates the chelation of extracellular calcium ions significantly mitigated the calcium effects of each drug. Likely, each drug under normal conditions promotes calcium entry via the plasma membrane. Because sigma receptors are known to localize at the plasma membrane, it is likely these compounds are activating those membrane sigma receptors, in turn opening channels to provide calcium entry into the cytosol. In terms of thapsigargin pretreatment, only siramesine resulted in a significant decrease in calcium levels. This demonstrates that under normal conditions, siramesine also promotes calcium flux from the endoplasmic reticulum into the cytosol.

Overall, this research entailed the use of sigma receptor ligands on BV2 microglia cells to study their effect on cytosolic calcium modulation. Importantly, we found a significant increase in intracellular calcium levels after 15 minutes of 1 µM treatment. Both sigma 1 and 2 receptor agonists have been shown to induce calcium increases, supporting our findings (Schmidt & Kruse, 2019). However, in contrast to other reports, we observed peaked cytosolic calcium after 15 minutes at 1 µM treatment, as opposed to 100 µM resulting in peak calcium levels after 1.5 – 3 minutes (Vilner and Bowen 2000). However, they used a variety of different sigma 1 receptor ligands and focused on endoplasmic reticulum calcium release in a different cell type as their cytosolic calcium increase occurred in the absence of extracellular calcium. In contrast, we took both extracellular and endoplasmic reticulum released calcium into account. A later study demonstrated the synthetic sigma 2 receptor agonist F281 induced a non-transient rise in cytosolic calcium from the endoplasmic reticulum and mitochondria (Cassano et al. 2009). However, this study used a much greater concentration of ligand (0.1 mM) compared to our 1 µM treatment. Nevertheless, their results corroborate our findings that sigma 2 receptor agonists can evoke cytosolic calcium increases through intracellular stores.

One question remaining is the connection between increased cytosolic calcium and microglia activation. Given the current literature, increased calcium can activate both pro and anti-inflammatory pathways in microglia. In terms of an anti-inflammatory route, one study demonstrated increased calcium flux from the endoplasmic reticulum to mitochondria can promote mitochondrial function, thereby inducing an anti-inflammatory microglial activation (Ooi et al. 2021). This increased calcium flow is stabilized by sigma receptors, suggesting the use of sigma receptor ligands in activation of a neuroprotective microglia. Potentially, sigma receptor ligands are activating the receptors, thereby inducing cytosolic calcium entry and increasing the uptake at the endoplasmic reticulum, in turn increasing flux at the mitochondria-associated membranes. This could influence more of a protective, anti-inflammatory microglia route of activation. A pro-inflammatory route could also occur due to the increases in calcium. Intracellular calcium is necessary for proper protein function, phosphorylation and kinase cascades, and gene transcription. All these actions are crucial for microglia activation, including cytokine and chemokine production, nitric oxide production, and phagocytic abilities. An increase in cytosolic calcium through sigma receptor ligands could potentially activate mechanisms that promote a pro-inflammatory phenotype.

## Data Availability Statement

The authors declare that all the data supporting the findings of this study are contained within the paper.

## Authorship Contributions

*Conceptualization, Funding acquisition, Project administration, Supervision, Visualization:* Simin, Lodholz, Schober

*Resources:* Schober, Crider, Neumann

*Methodology, Investigation, and Formal analysis:* Lodholz, Simin, Schober

*Writing, original draft:* Simin, Lodholz, Schober

*Writing, review & editing:* Simin, Schober

